# Hyphal induction in *Candida albicans*: Optimizing medium parameters to accelerate hyphal growth enhanced by GlcNAc

**DOI:** 10.1101/2025.06.12.659427

**Authors:** A Rachana, PV Atheena, Allan Britto J, Uttara Chakraborty, Ritu Raval

**Affiliations:** Department of Biotechnology, Manipal Institute of Technology, Manipal Academy of Higher Education, Manipal, Karnataka-576104, India; Manipal Institute of Regenerative Medicine, BSF Campus, Yelahanka, Govindapura, Manipal Academy of Higher Education, Bengaluru, Karnataka-560064, India

**Keywords:** *Candida albicans*, Candidiasis, GlcNAc, Hyphal induction, Medium optimization.

## Abstract

Hyphal formation is a critical virulence trait in *Candida* species, contributing significantly to host tissue invasion, immune evasion, and disease progression. The morphological transition from yeast to hyphal form is therefore a key target for antifungal intervention. However, conventional hyphal induction media are often complex, expensive, and poorly standardized. In this study, we developed a simplified and cost-effective medium (MF8) containing peptone (0.16%), dextrose (0.4%), and bovine serum albumin (BSA; 0.25%), which supported moderate hyphal growth under nutrient-limited conditions. The addition of N-acetylglucosamine (GlcNAc) markedly enhanced filamentation. Using a Design of Experiments (DOE) approach via JMP software, we evaluated the effects of GlcNAc and magnesium sulfate (MgSO₄) on hyphal induction. GlcNAc was identified as a significant enhancer (p < 0.05; R² = 0.26), while MgSO₄ had no significant impact (p > 0.05). Under optimized conditions (30 mM GlcNAc, 1 mM MgSO₄), RT-qPCR analysis revealed strong upregulation of *HWP1* (100-fold at 4 h; 85-fold at 6 h) and *HGC1* (>6-fold; p < 0.05). A concurrent pH shift from alkaline to acidic during 1–6 hours correlated with hyphal induction and gene activation, suggesting that acidification may serve as an additional cue regulating morphogenesis.

**IMPORTANCE:** *Candida* species are highly adaptable opportunistic fungi that persist in diverse host niches due to numerous virulence factors. While typically commensal, they can transition to pathogenic forms, with hyphal formation being a key virulence trait. This yeast-to-hypha switch facilitates tissue invasion, immune evasion, and disease progression, making it a major target in antifungal research. Conventional hyphal induction media are often complex and costly. Developing a simplified, cost-effective, and reproducible medium would enhance studies on morphogenesis and pathogenesis, support drug screening, and potentially reveal novel virulence mechanisms under alternative environmental cues.

## INTRODUCTION

A diverse array of microbes significantly impacts human health, and the diseases caused due to fungi remain often overlooked despite the vast Kingdom of Fungi encompassing over 2 million species. Certain fungi lead to severe health consequences, contributing to approximately 2 million deaths annually (4, 26). Candida species, cause majority of the fungal infections, particularly *Candida albicans,* being predominant in causing candidiasis infections. The World Health Organization (WHO’s) fungal pathogen priority list categorizes both *C. albicans* and *C. auris* as critically important pathogens, further underscoring the severity of their impact (13, 17). *Candida albicans* is an opportunistic pathogen that exists as a commensal organism in 50 % of healthy individuals, colonizing the mucosal membranes of the oral cavity, gastrointestinal tract, and vaginal area. Various factors contribute to its switch from a commensal to a pathogenic state owing to the disturbances in normal homeostasis, resulting in clinical infections that range from superficial mucocutaneous infections to severe systemic infections. This polymorphic fungus can exist in either a budding yeast form or a filamentous hyphal and pseudohyphal form. The pseudohyphal form is characterized by highly branched ellipse with distinct constrictions at cellular junctions, whereas the hyphal form exhibits true septa without constrictions, parallel-sided walls, and relatively less branching. Numerous studies have demonstrated that the ability to reversibly switch between these morphogenic states plays a significant role in virulence and the manifestation of infection (8,27). This hyphal form of *Candida* evades the immune system by piercing and killing macrophages, as well as penetrating host cells through cytoplasm-driven forces resulting from turgor pressure. Additionally, it secretes a toxin called candidalysin, which damages epithelial cells and facilitates bloodstream infections (2,6,14). This invasive nature of the hyphal form is a critical virulence trait, making it a significant target for therapeutic interventions against infections. The morphological switch from yeast to hyphal is caused by several environmental cues, including temperature, pH, N-acetyl glucosamine (GlcNAc), CO_2,_ and serum (3,19). To trigger the yeast-to-hyphal transition *in-vitro*, research groups have often employed synthetic medium. One of the most utilized options is Lee’s synthetic medium, which facilitates the development of either pure yeast or pure hyphal cells. The germ tube formation is typically induced within 3 to 4 hours. However, to achieve a fully developed mycelial form, the cultures are recommended to have an incubation time of 18 to 27 hours. Lee’s medium comprises 14 components, including 8 amino acids, 4 inorganic salts, biotin, and glucose, making it a complex formulation. These components, being expensive, also add to the cost of the medium (11). Another well-known inducer of hyphal formation is GlcNAc. The addition of 2.5 mM GlcNAc to a culture of pure blastospores, derived from the late logarithmic phase, at pH 6.6 and a temperature of 37 °C, leads to germ tube formation within 5 hours. Supplementing the medium further with 0.1 mM of MgSO_4_ or MnCl_2_ enhances germ tube formation, with many cells developing into true hyphae. However, an increase in concentration beyond this leads to the overall reduction in the germ tube germination (24). Another supplement that has been used in the experimental set-up for hyphal induction is serum, but its use is impeded due to its high cost (18,25). In another study, research group used spider medium, GlcNAc, and RPMI 1640 medium to induce hyphal growth, the hyphal induction could be induced after 48 hours (10). The medium used for inducing the yeast-to-hyphal transition contain either complex and expensive components and long experimentation incubations periods.

The aim of the study was to optimize a medium with simple and cost-effective medium components for hyphal induction and further evaluating the effects of nutrient-limited conditions in combination with known hyphal inducers, such as N-acetylglucosamine (GlcNAc), and selected inorganic salts on hyphal induction using a Design of Experiments (DOE) approach via JMP software. While GlcNAc is known to promote hyphal formation, its effectiveness was variable under nutrient limited conditions (MF8) failing to induce filamentation in certain combinations, while successfully triggering it in others. The resulting optimized medium not only provides a simplified platform for investigating the yeast to hyphal transition, but also holds potential for antifungal drug screening, particularly for compounds targeting morphogenetic switching. Moreover, insights gained from the conditions promoting this transition may further provide insights into the regulatory mechanisms underlying *Candida*’s morphological plasticity and pathogenesis.

## MATERIALS AND METHODS

### Strain and Culture Conditions of *C. albicans*

The *Candida albicans* strain ATCC 10231 was used in this study. For routine cultivation, the organism was maintained on Sabouraud Dextrose Agar (SDA) plates, which were incubated at 37 °C and subsequently stored at 4 °C. For long term preservation, strains were stored in 30% glycerol stocks at -20 °C. Prior to experimental use, *C.albicans* cells were revived from glycerol stocks by streaking onto fresh SDA plates, followed by overnight pre-cultivation in Sabouraud Dextrose Broth (SDB) at 37 °C with constant agitation at 180 rpm. To induce hyphal formation, pre-cultures were inoculated into specific medium modifications optimized for hyphal induction.

### Development of optimized medium for hyphal induction

To induce rapid hyphal formation, three key carbon and nitrogen sources—dextrose, peptone, and bovine serum albumin (BSA), respectively—were initially modified to limit their concentrations in the medium. This adjustment was made based on the observation that *Candida* cells, under low glucose or nutrient-deprived conditions, tend to utilize amino acids as alternative carbon sources (3). Nutrient stress may trigger a survival response in *Candida*, promoting hyphal formation. Various medium compositions with differing proportions of carbon and nitrogen sources were analysed. The medium that successfully induced hyphal development were then enriched with additional components such as GlcNAc, MgSO_4_, and MnCl_2_. The effects of these additives on hyphal induction, mycelial colony formation, and the rate of formation were subsequently observed.

### Optimization of nutrient limited medium

Two standard fungal growth medium, Broth SDB Broth and Yeast Peptone Dextrose Broth (YPD), were utilized to observe the hyphal induction in *Candida albicans.* Observations were conducted using light microscopy at 40 X and 100 X magnifications over a 40 hours period, with every two hours sample collection interval. Subsequently, modifications were made to the YPD Broth, substituting yeast extract with bovine serum albumin (BSA) Fraction V, while maintaining bacteriological peptone at 20 g/L (2 %) and dextrose at 20 g/L (2 %). A total of ten modifications were performed, adjusting the concentrations of peptone, dextrose, and BSA, varying between 0.1 % to 2 %. The impact of these modifications on morphogenic transitions were assessed using light microscopy at both 40 X and 100 X magnifications over an 8 hours period, with every 1 hour sample collection intervals. The filling volume was maintained at 24 % with temperature at 37 °C and an agitation speed of 200 rpm, using 1 % inoculum for all experimental conditions.

### Supplementation of optimized nutrient limited medium

The optimized nutrient-limited medium, which demonstrated hyphal development, was further supplemented with GlcNAc, MgSO_4_ and MnCl_2_ to enhance the hyphal formation. In initial experiments, GlcNAc in the medium was at concentrations of 30 mM, 50 mM, and 100 mM. Subsequently, MgSO_4_ was evaluated in combination with GlcNAc at concentration ranging from 0.05 mM to 1 mM and 2.5 mM to 30 mM respectively. Finally, MnCl_2_ was assessed in conjunction with GlcNAc and MgSO_4_ at concentrations between 5 mM and 20 mM. All experiments were conducted with a 24 % filling volume, at a temperature of 37 °C, and an agitation speed of 200 rpm, using 1 % inoculum. The different combinations of these components were generated using JMP software (https://www.jmp.com/en/software), customized DOE (Design of Experiments). The impact of these combinations on mycelial development was observed through light microscopy at 40 X and 100 X magnifications.

### Customized DOE (Design of Experiment) and Statistical Analysis or Experimental Design

To design the experiment, statistical tool JMP software® (Version 18, SAS Institute, USA) was imposed. Since the parameters were discrete numeric values with varying intervals between them, custom design was chosen to analyse the impact of parameters on the responses. The selected independent variables were concentration of GlcNAc and MgSO_4_ at four different levels. Hyphal induction was the only dependent variable. All dependent and independent variables are presented in Table 2 along the experimental design including the levels of GlcNAc and MgSO_4_. The fit model option was used to analyse the multiple independent variables (predictors) and their relationship with a dependent variable (response). Analysis of variance (ANOVA) was used to determine if there are statistically significant differences between the means of different groups. JMP prediction profiler was used to find optimal settings for individual parameters.

### Total RNA extraction and Quantitative real time PCR (qRTPCR) analysis

For the analysis of gene expression, total RNA was extracted using acid phenol method as previously described (1). Hyphal cells cultured in optimized hyphal inducing medium (Run 20) were pelleted from different time points (1 hour, 2 hours, 4 hours and 6 hours) by centrifugation at 10,000 rpm, 10 °C, 10 mins (Thermo Scientific Multifuge 1S-R Table Top Centrifuge), the cells were resuspended in 270 µL of sodium acetate buffer (Sodium acetate, 50 mM, pH 7 and EDTA, 10mM, pH 8) and 30 µL of 10 % SDS was added. Glass beads (0.5 mm) were added equivalent to the size of the cell pellet, followed by addition of 300 µL of acid phenol preheated at 65 °C, the entire mixture was vortexed for 1 min and then incubated at 65 °C for 30 mins and then 300 µL of chloroform was added and centrifuged at 10,000 rpm for 10 mins at 4 °C. The aqueous phase is collected separate from the organic phase and 1/10^th^ volume of 3M of sodium acetate is added to the aqueous phase and then 2.5 times of chilled absolute ethanol is added and incubated at -20 °C for 2 hours and later centrifuged at 13,000 rpms for 10 mins at 4 °C (Sorvall™ Legend™ Micro 21R Microcentrifuge). The extracted RNA pellet was air dried and washed twice using 70 % ethanol, and the air-dried pellet was dissolved using molecular grade water. The RNA was quantified using nanodrop instrument (Nanodrop One, Thermofischer Scientific) at 260 nm. The RNA was transcribed into cDNA using PrimeScript™ RT Reagent Kit (Takara, catalogue no. RR037A). The quantitative PCR was performed using TB green® Premix Ex Taq™ II (Takara, catalogue no. RR820A) on QuantStudio 5 applied biosystems (Thermo Fischer Scientific). *ACT1* gene was used as the endogenous control for normalization of gene expression. Primers used for quantitative real time polymerase chain reaction (qRTPCR) of *HWP1* and *HCG1* are listed in Table 5. A one-way ANOVA was performed using GraphPad PRISM software version 10.4.1, followed by Bonferroni test to analyse the significance in relative fold change between control and multiple time points.

### Statistical Analysis

Data for gene expression analysis was expressed one-way ANOVA (Analysis of Variance) performed using GraphPad PRISM software version 10.4.1, followed by Bonferroni test to analyse the significance in relative fold change between control and multiple time points. A value of (*p* < 0.05) was considered statistically significant.

## RESULTS AND DISCUSSION

### Effect of Nutrient Limitation

Nutrients are essential components required for the survival, reproduction, and development of microbes. The ability to sense and respond to the nutrient availability enables them to acquire nutrients and survive from various ecological niches (23). One of the virulence factors of *C. albicans* is its capacity to survive and adapt to rapidly changing environments, diverse host niches, and stress conditions such as elevated temperature, high oxidative stress, and nutrient limitation (2,16). This study investigated the effect of nutrient limitation on the yeast to hyphal transition in *C.albicans.* Observations over a 40 hours period in nutrient rich medium, specifically SDB and YPD Broth, revealed the presence of a few hyphal cells at 4 hours and 6 hours, respectively. However, beyond these time points, no hyphal cells were observed. The subsequent experiments were conducted over an 8 hours period, focusing on the early hours where hyphal formation was evident. YPD Broth was modified by replacing yeast extract with bovine serum albumin (BSA), resulting in a total of 10 variations (**Table 1**).

**Table 1.** Experimental variants of Yeast Peptone Dextrose (YPD) medium formulated by replacing yeast extract with bovine serum albumin (BSA) to evaluate its efficacy as an alternative complex nitrogen source.

| Media<br>Formulation (MF) | Peptone % | Dextrose<br>% | BSA<br>% |
| --- | --- | --- | --- |
| MF 1 | 0.5 | 0.5 | 1.0 |
| MF 2 | - | 0.1 | 0.5 |
| MF 3 | 1.0 | 0.5 | 1.0 |
| MF 4 | 0.5 | 0.5 | 2.0 |
| MF 5 | 0.5 | 0.2 | 1.0 |
| MF 6* | 0.5 | 0.5 | 0.5 |
| MF 7* | 0.2 | 0.5 | 0.5 |
| MF 8 | 0.16 | 0.4 | 0.25 |
| MF 9 | 0.1 | 0.5 | 0.2 |
| MF10 | 0.1 | 0.5 | 0.1 |
| * Media formulations were tested at pH 6.0 and 7.4 |  |  |  |

The original YPD medium comprises of 2 % peptone and dextrose, and 1 % yeast extract. Initially, two modifications (MF1 and MF2) were implemented. In MF1, peptone and dextrose concentrations were reduced to 0.5 %, while BSA was maintained at 1 %, equivalent to the original concentration of yeast extract. Under these conditions, germ tube formation was observed at 2 hours, with distinct hyphal structures appearing from 4 to 8 hours, although yeast cells were also present. In MF2, peptone was entirely omitted, and dextrose and BSA were reduced to 0.1 % and 0.5 %, respectively. The exclusion of peptone in formulation MF2 completely abrogated hyphal formation and significantly impaired yeast proliferation, as indicated by reduced optical density (OD₆₀₀). To delineate the contribution of individual medium components, MF3–MF5 were derived from MF1 by selectively altering peptone (increased to 1% in MF3), BSA (increased to 2% in MF4), and dextrose (reduced to 0.2% in MF5), while keeping other parameters constant. Despite these modifications, only limited hyphal structures were observed, suggesting that elevated nutrient concentrations do not favour morphogenesis. MF6 and MF7, formulated with reduced BSA (0.5%) and combined reductions in BSA (0.5%) and peptone (0.2%), were evaluated at pH 6 and 7.4 to investigate pH-dependent effects. Hyphae appeared by 3 hours, but no substantial enhancement was noted. Further reductions in MF8–MF10 were tested to refine the nutrient profile. While MF9 and MF10 (BSA and peptone ≤0.2%) elicited weak responses, MF8 (BSA 0.25%, peptone 0.16%, dextrose 0.4%) yielded pronounced hyphal development from 4 to 8 hours, along with visible mycelial colonies at later time points (7–8 hours, **Fig. 1**). These results underscore the importance of carefully optimized nutrient limitation in promoting *Candida albicans* hyphal differentiation under defined *in-vitro* conditions.

**Fig. 1.**
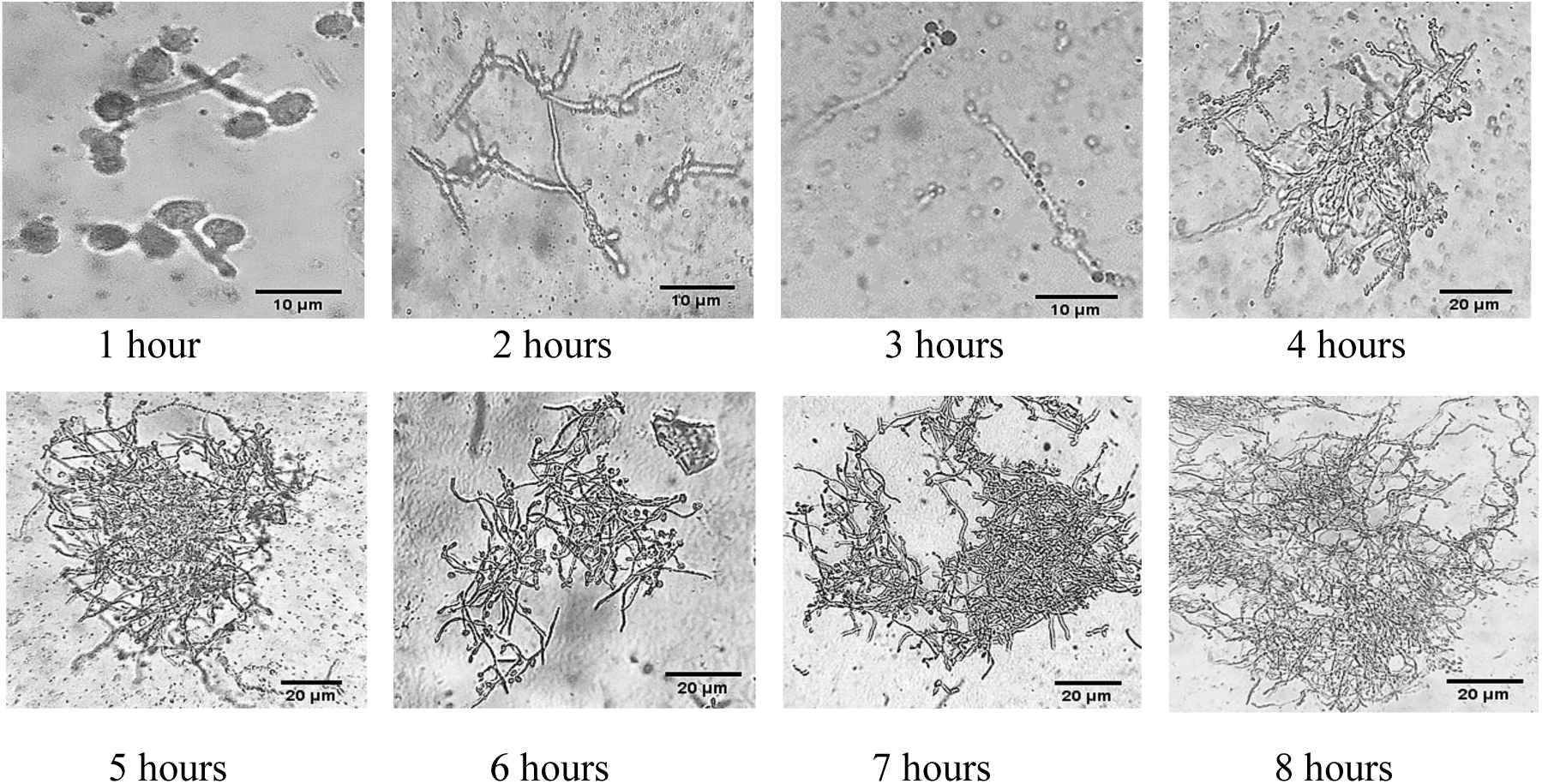
Hyphal formation observed in Media Formulation 8 (MF8), which comprises of 0.16 % peptone, 0.4 % dextrose, and 0.25 % BSA. Mycelial colonies were observed at the 7 and 8 hours’ ime points. Representative microscopic slides with *n*=3

### Effect of Supplementation in Nutrient Limited Medium

Medium Formulation 8 (MF8) was further refined to enhance and accelerate hyphal development by supplementing with GlcNAc, MgSO_4_, and MnCl_2_. Initially, the influence of GlcNAc as a single factor was explored at concentrations of 30 mM, 50 mM, and 100 mM in the MF8 medium. Both 50 mM and 100 mM concentrations were observed with hyphal induction and clumped mycelial growth, as shown in **Fig. 2**, with germ tube formation observed at the 1 hour time point. Conversely, the 30 mM concentration did not induce hyphal formation. Subsequently, the effects of combining 30 mM GlcNAc with varying concentrations of MgSO_4_ (0.05 mM, 0.1 mM, 0.25 mM, and 1 mM) were investigated (Table 2). Additionally, other GlcNAc concentrations (2.5 mM, 5 mM, 10 mM, and 25 mM) were tested in combination with MgSO_4_ using a design of experiments approach facilitated by JMP software. Among the combinations generated, levels 5, 7, 9, 10, 11, 12, 13, 17, 18, 19, and 20 successfully induced hyphal formation, as depicted in **Fig. 3**. Furthermore, the combination of manganese and GlcNAc was evaluated under nutrient limited conditions for hyphal induction, but no hyphal formation was observed.

**Fig. 2.**
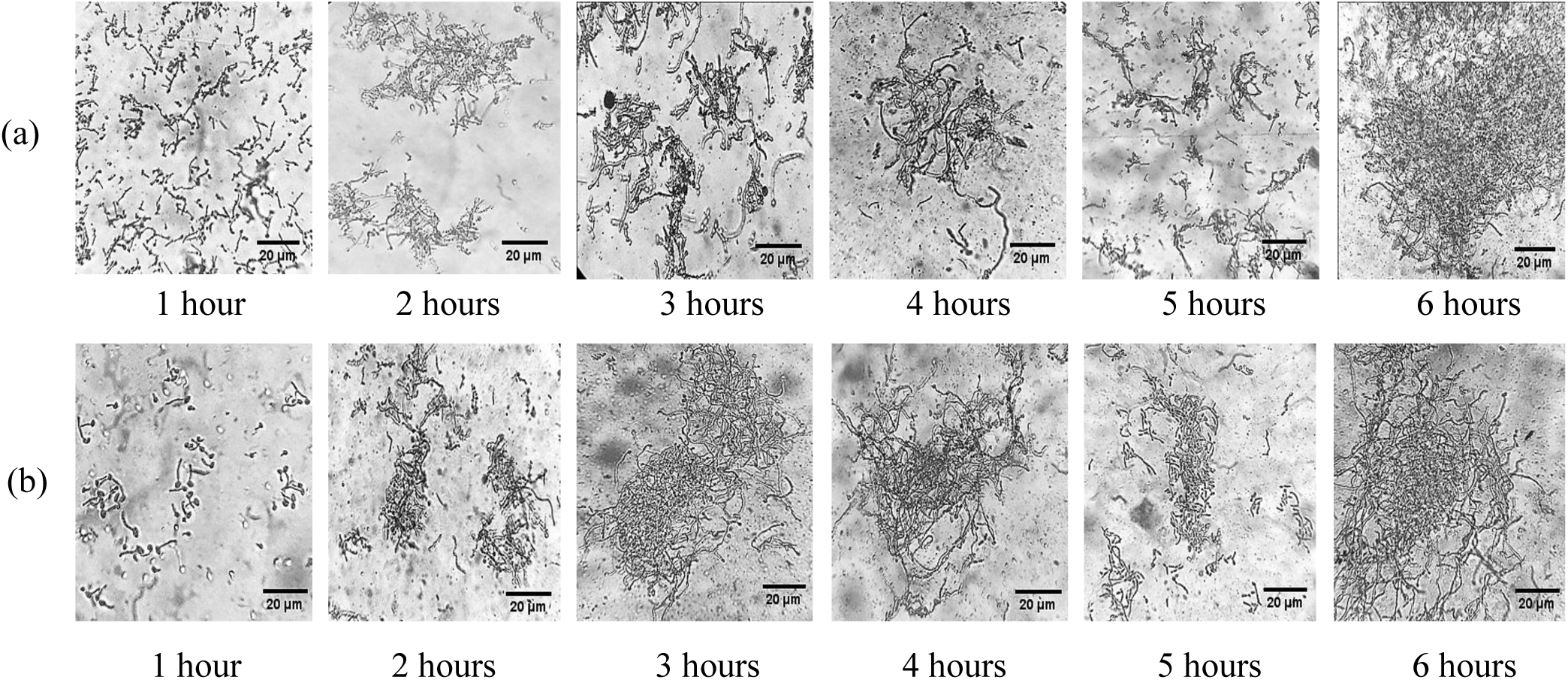
Hyphal formation observed in (a) Media Formulation 8 (MF8) supplemented with 30 mM GlcNAc and (b) Media Formulation 8 (MF8) supplemented with 100mM GlcNAc. Representative microscopic slides with *n*=3

**Fig. 3.**
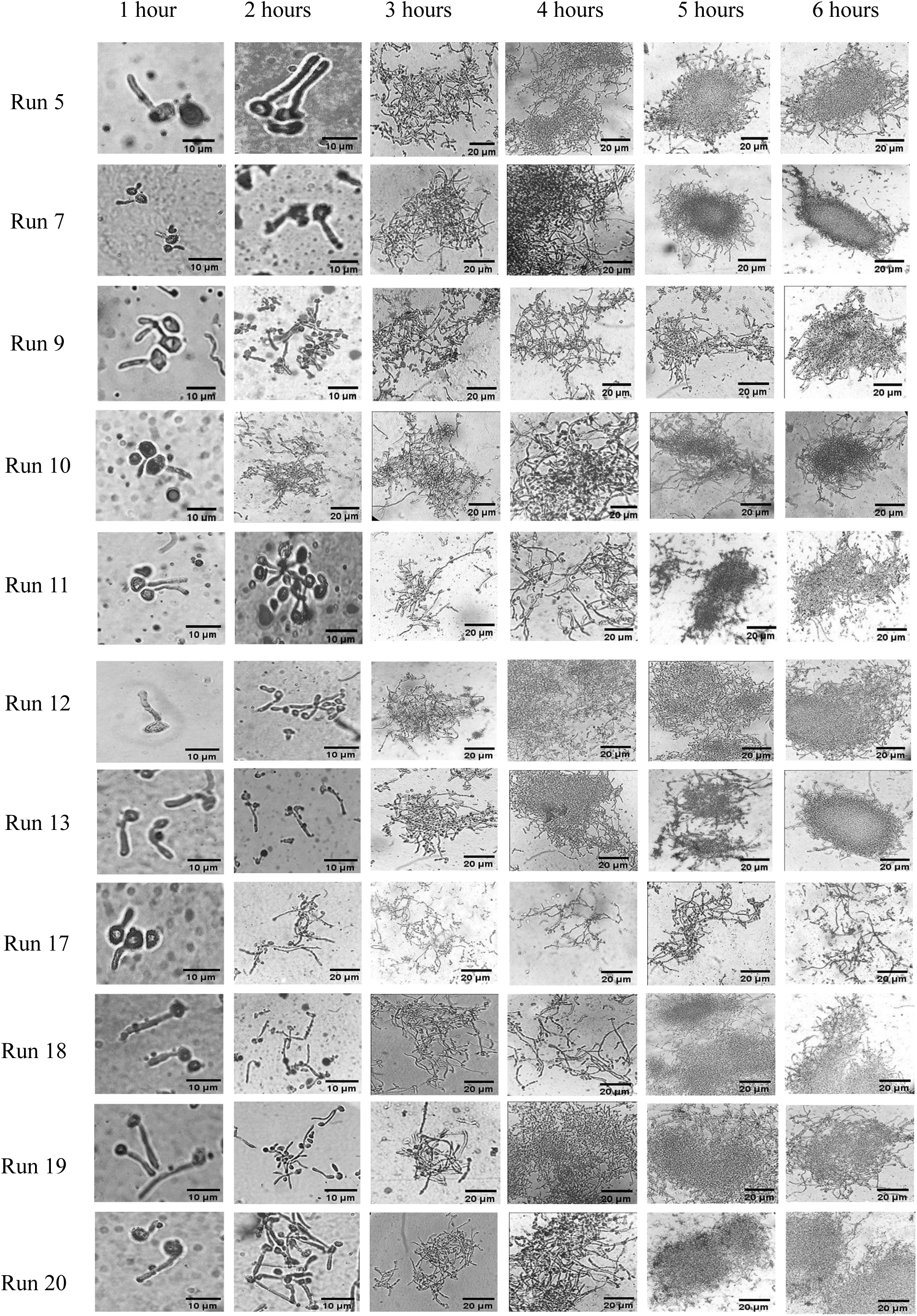
Hyphal formation and mycelial growth observed in different combination runs of GlcNAc and MgSO_4_. Representative microscopic slides with *n*=3

**Table 2.**
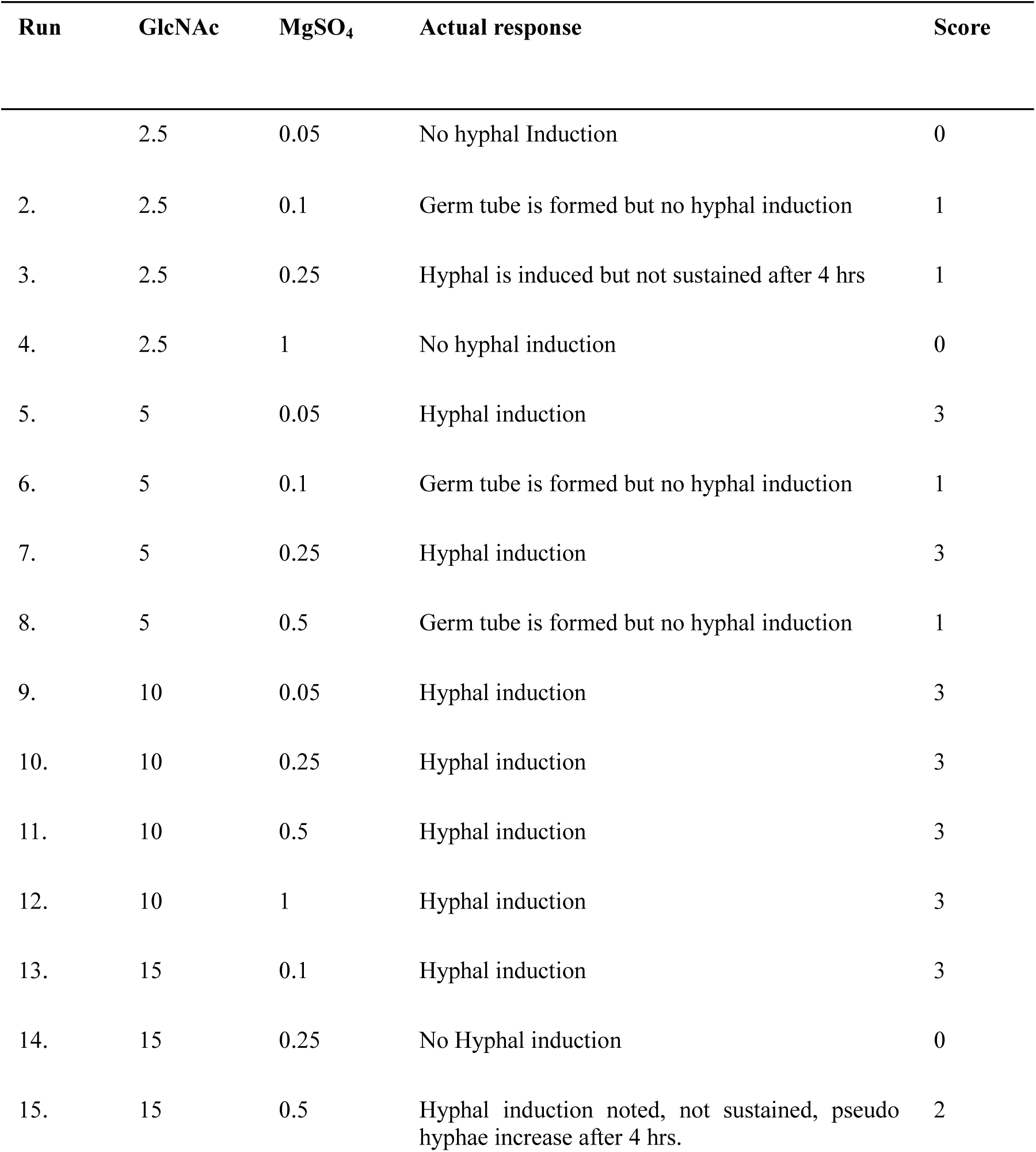
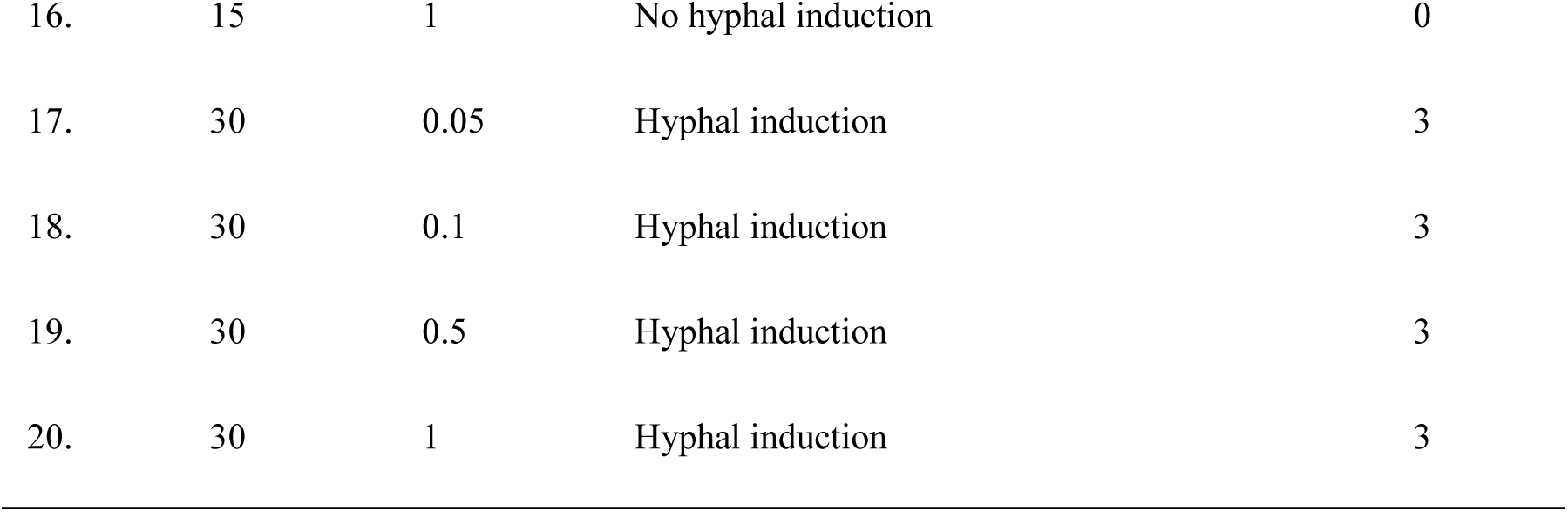
Combinatorial treatments of N-acetylglucosamine (GlcNAc) and magnesium sulfate (MgSO₄) at varying concentrations were designed using JMP software, yielding a total of 20 experimental runs along with their corresponding response data.

The amide derivative sugar GlcNAc (N-acetylglucosamine) is an important component of the cell wall of fungi composed of polysaccharide chitin. GlcNAc has multiple roles in metabolism and import, scavenging, yeast to hyphal transition and GlcNAc induced cell death. It serves as a nutrient source for many microbes, including *Candida albicans,* and is known to be a potent inducer of hyphal development (22). GlcNAc uptake in *Candida albicans* activates both catabolic and anabolic gene expression pathways. Transport into the cell is mediated by Ngt1, followed by phosphorylation to GlcNAc-6-phosphate via Hxk1. In the anabolic pathway, GlcNAc-6-phosphate is converted to UDP-GlcNAc, an essential precursor for chitin biosynthesis—a major structural component of the fungal cell wall, particularly concentrated at the hyphal tip during polarized growth (15, 20). Chitin deposition at these sites is catalyzed by distinct classes of chitin synthases. Liss and Slater (12) first identified GlcNAc as a potent inducer of germ tube formation and reported enhanced filamentation in the presence of MgSO₄ and MnCl₂. In this study, GlcNAc was employed to promote hyphal induction under nutrient-limiting conditions. Metals such as manganese are essential micronutrients that play key roles in microbial physiology and virulence. In *C. albicans*, manganese uptake is mediated by Smf12 and Smf13 transporters; deletion of both impairs hyphal development and virulence (28). However, in the present study, the addition of manganese in combination with GlcNAc did not enhance hyphal formation under MF8 conditions, contrary to earlier findings. This suggests that nutrient context critically influences metal ion-mediated modulation of morphogenesis.

### Experimental Design Analysis

The optimization of medium using statistical tools ensures optimal balance between input variables and output responses in experimental designs, enhancing the overall efficiency and effectiveness of the medium optimization process. The selection of a specific design intensifies the generation of an optimal collection of experimental designs to examine the influence of independent factors on the selected responses. In the present study, custom design was used to develop a conceptual framework for medium optimization and thereby enabling to explore the design space using graphical and mathematical design. In **Table 3**, it is evident that the method is influenced only by the factor concentration of GlcNAc from the *p* value (*p* <0.05).

**Table 3.**
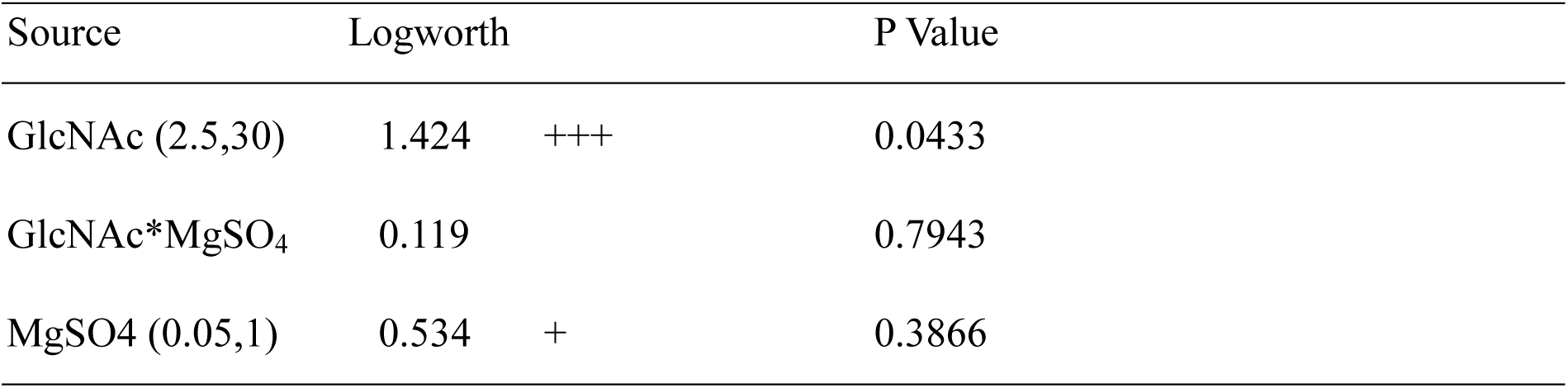
Effect summary of custom design.

| Source | Logworth | P Value |
| --- | --- | --- |
| GlcNAc (2.5,30) | 1.424 +++ | 0.0433 |
| GlcNAc*MgSO<sub>4</sub> | 0.119 | 0.7943 |
| MgSO<sub>4</sub> (0.05,1) | 0.534 + | 0.3866 |

### Model fit analysis

To assess the adequacy and predictive strength of the experimental model, data from all design runs were incorporated into the statistical analysis. As depicted in **Fig. 4**, the concentration of N-acetylglucosamine (GlcNAc) emerged as the only statistically significant factor influencing hyphal induction, with a p-value of 0.043 (p < 0.05) and a corresponding coefficient of determination (R²) of 0.26. Although the R² value is relatively low, this is largely attributable to the qualitative nature of the response variable. Hyphal induction was evaluated based on a semi-quantitative morphological scoring system ranging from 0 to 3, where: 0 indicates no hyphal formation; 1 denotes germ tube initiation without true hyphal elongation; 2 represents partial or unsustained hyphal development; and 3 corresponds to robust and sustained hyphal formation. This categorical scale was designed to capture gradations in morphological transition under different experimental conditions but does not offer the resolution of a continuous variable, thereby limiting the model’s overall fit.

**Fig. 4.**
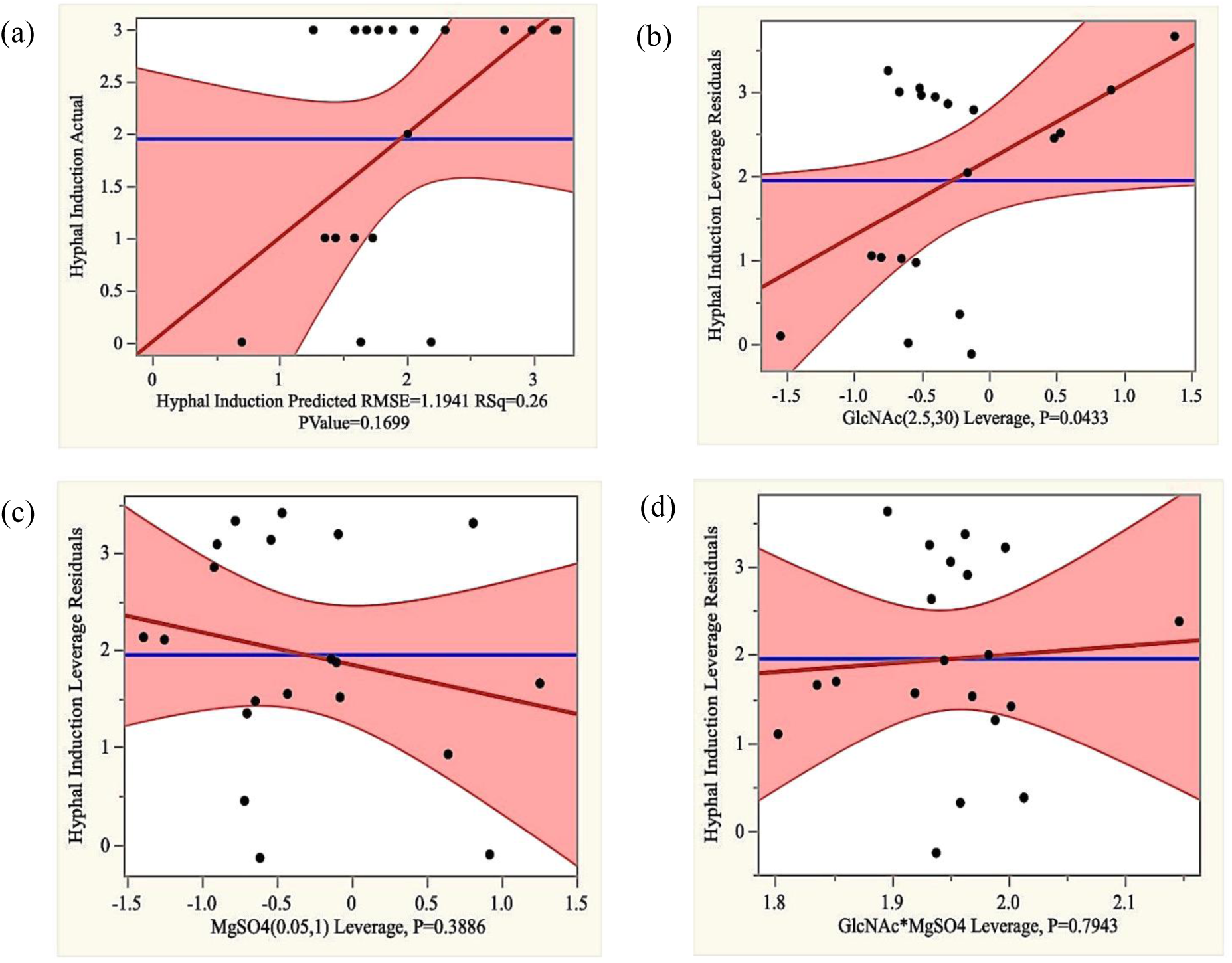
Multiple linear regression analysis, visualized via JMP leverage plots, revealed a statistically significant correlation (*p* < 0.05) between GlcNAc concentration and hyphal induction, while MgSO_4_ concentration and the GlcNAc-MgSO_4_ interaction showed no significant effects (*p* > 0.05) (a) Actual v/s predicted plot for selected responses, (b)leverage plot of GlcNAc against hyphal induction (c) leverage plot of MgSO_4_ against hyphal induction (d) leverage plot of interaction of GlcNAc and MgSO_4_ against hyphal induction.

Nonetheless, the analysis clearly indicates that GlcNAc concentration plays a critical role in promoting the yeast-to-hypha transition, whereas other tested variables, such as MgSO₄, did not show statistically significant

### Statistical Analysis for Hyphal Induction

The model yielded an F-value of 5.13, indicating statistical significance for the GlcNAc variable within the design space. According to this analysis, there is only a 3.7% probability that such a high F-value could result from random noise, thereby supporting the model’s reliability in identifying significant factors. A summary of the statistical analysis is presented in **Table 4**. From these results, it is evident that GlcNAc concentration significantly contributes to hyphal induction (p < 0.05), whereas MgSO₄ did not exert a statistically significant effect on the response (p > 0.05), underscoring its limited role under the experimental conditions evaluated.

**Table 4.** Summary of the statistical analysis for hyphal induction.

| Source | Sum of squares | DF | F-value | p-value | Remarks |
| --- | --- | --- | --- | --- | --- |
| Model | 7.18 | 3 | 2.11 | 0.169 |  |
| GlcNAc | 6.19 | 1 | 5.13 | 0.043 | Significant |
| MgSO<sub>4</sub> | 1.42 | 1 | 1.184 | 0.386 | Non-significant |
| GlcNAc* MgSO<sub>4</sub> | 0.11 | 1 | 0.096 | 0.794 | Non-significant |
| R<sup>2</sup> | R<sup>2</sup>(adj) |  |  |  |  |
| 28% | 15% |  |  |  |  |

**Table 5.**
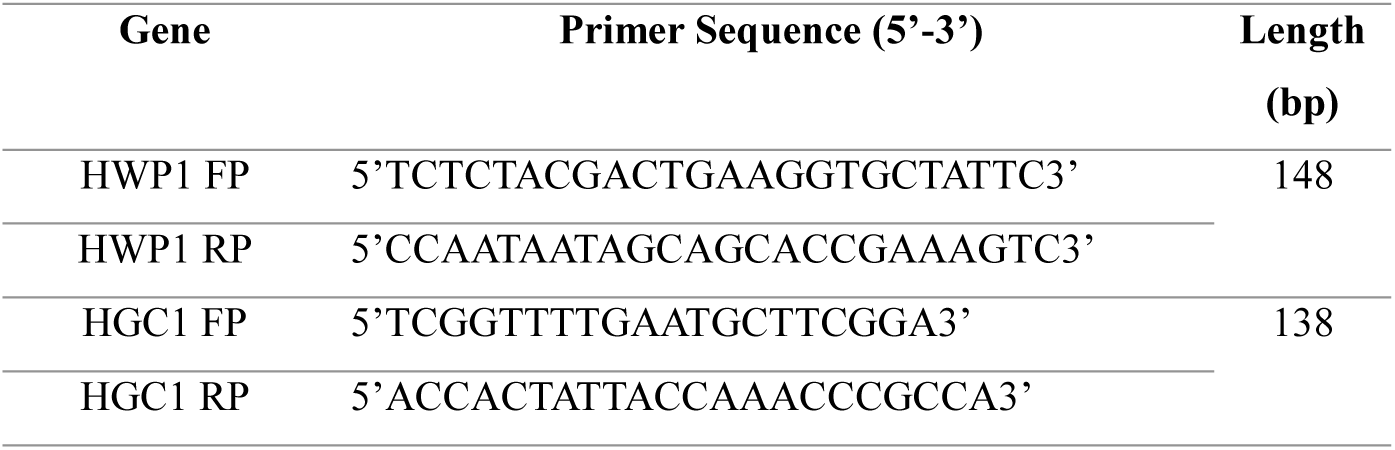
List of primers for hyphal-specific genes used in qRTPCR analysis.

### Statistical Optimization Using Prediction Profiler

The prediction profiler was utilized to visualize the continuous relationship between input variables (GlcNAc and MgSO₄ concentrations) and the phenotypic response of hyphal induction. This tool enables exploration of parameter interactions and response trends across the tested concentration ranges. As illustrated in **Fig. 5**, an increase in GlcNAc concentration correlates positively with hyphal induction, reinforcing its role as a key morphogenetic trigger. In contrast, MgSO₄ did not exhibit a similar trend; in fact, higher concentrations were associated with a marginal decline in hyphal induction, suggesting a possible inhibitory effect at elevated levels. By applying the desirability function optimization within the profiler, the ideal condition for maximal hyphal induction was identified as 30 mM GlcNAc and 0.05 mM MgSO₄. These optimized parameters provide a refined framework for inducing robust hyphal development under nutrient-limited conditions, thereby facilitating future studies on morphogenesis and antifungal target screening.

**Fig. 5.**
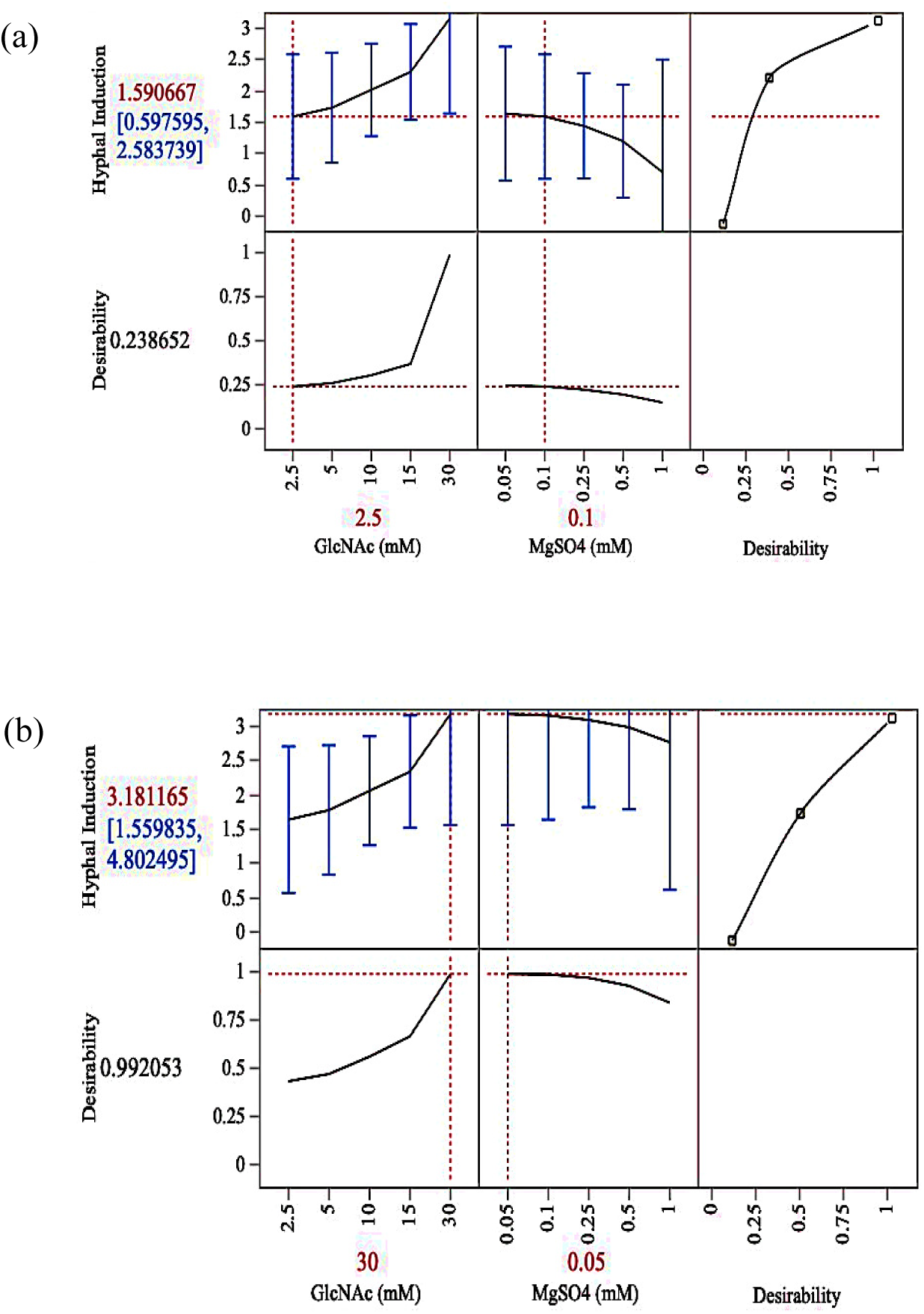
Desirability plot for the optimization of process parameters (a) plots before optimization and (b) plots after optimization.

### Gene expression analysis of hyphal transition in nutrient limited medium supplemented with GlcNAc

We analysed gene expression of key hyphal-specific genes, (Hyphal G1 Cyclin 1) *HWP1* and Hypha-specific G1 cyclin 1 (*HGC1)*, which are well-established markers of hyphal development in *C.albicans* under Run 20 (30 mM GlcNAc, 1 mM MgSO_4_) conditions using qRTPCR at different time points (1 hour, 2 hours, 4 hours and 6 hours). Morphological observations had previously confirmed hyphal induction at 4 hours and 6 hours, with germ tube formation at 1 hour and 2 hours. The *HWP1* gene encodes for hyphal wall protein 1, an adhesin crucial for host cell attachment, expressed on the hyphal surface in *C. albicans* and *HGC1* encodes a cyclin protein essential for the regulation of hyphal extension and polarized growth (5,9). qRTPCR analysis revealed significant upregulation of both genes under Run 20 conditions (Fig. 6). Notably, *HWP1* exhibited 50-fold increase in expression at the 1-hour time point compared to the control (*p* < 0.05), with a further increase to 100-fold and 85-fold at 4 and 6 hours, respectively (*p* < 0.05) (Fig. 6a). Similarly, *HGC1* expression was significantly upregulated at 2, 4, and 6 hours, showing relative fold change of 5.91, 7.09, and 6.07, respectively (*p* < 0.05) (Fig. 6b). We further analysed the pH during hyphal formation, interestingly, there was shift to alkaline pH at 1 hour and 2 hours and lowers to acidic pH during hyphal formation with 3.95 at 4 hours, further reduced to 2.26 at 6 hours time point (Supplementary Fig. 1) While alkaline pH has been reported to play a role in hyphal morphogenesis regulated by the Rim101 pathway (21), our study aligns with a previous study demonstrating that Candida can undergo hyphal morphogenesis even under acidic pH conditions. Notably, this earlier study also reported upregulation of hyphal specific genes like *HWP1* and *HGC1* under acidic pH regulated by Rfg1-Bcr1 mediumted pathway (7).

**Fig. 6.**
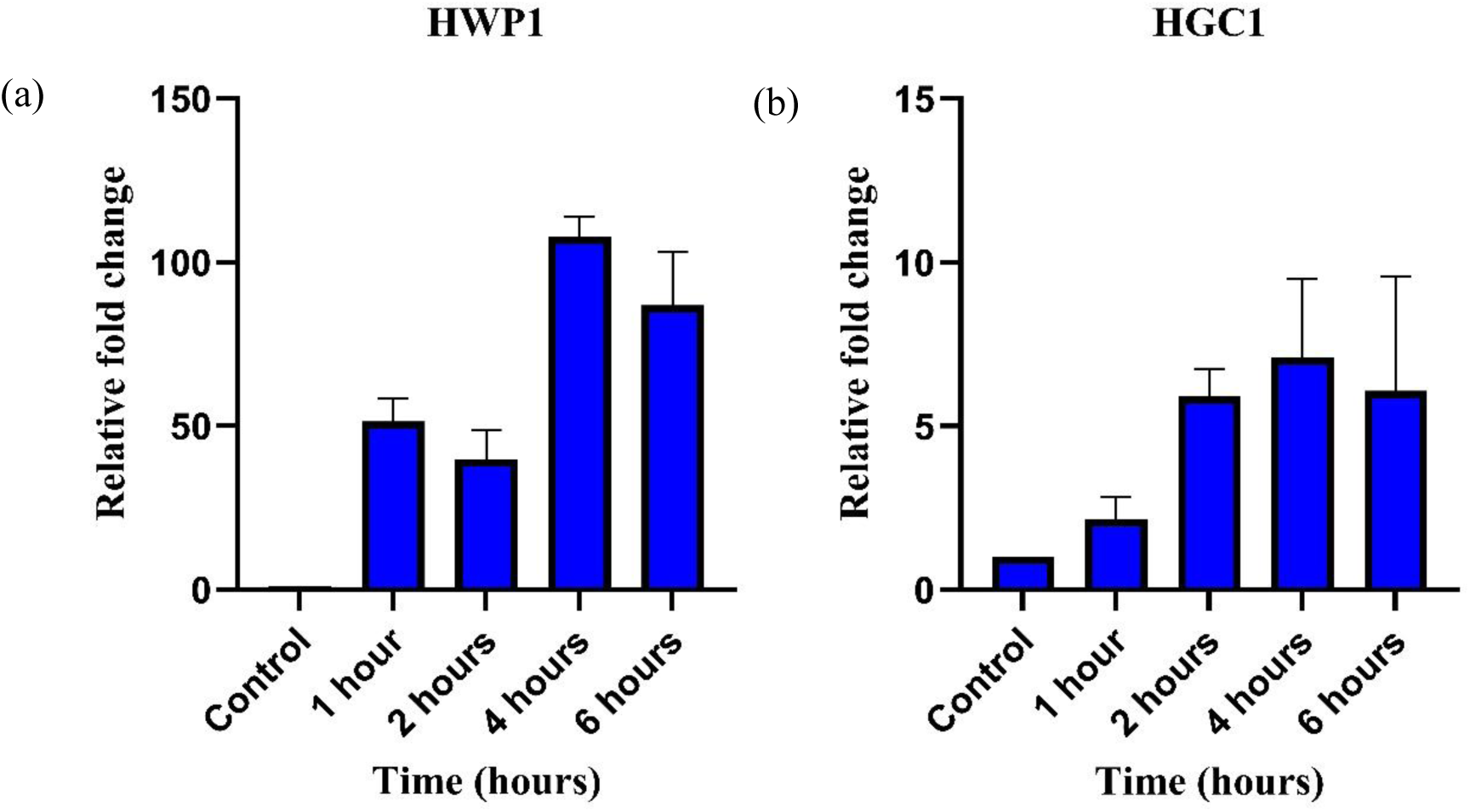
Gene expression analysis of hyphal-specific genes (a) Graph illustrating significant increase in the relative fold change of *HWP1* gene expression comparing the control to various time points corresponding to hyphal induction, a 50 fold increase at 1 hour and 100 and 85 fold increase at 4 hours and 6 hours respectively is observed (b) significant increase in relative fold change of *HGC1* gene with 5.91, 7.09, and 6.07 fold change is observed for 2 hours, 4 hours and 6 hours. All data are presented as the mean ± SD, *n*=3; compared to the control.

## CONCLUSION

In this study, the YPD medium was modified and optimized for inducing hyphal formation, MF8 medium containing 0.16 % peptone, 0.4 % dextrose, and 0.25 % BSA, induced moderate hyphal formation between the 4 hours and 8 hours time points. Supplementation with higher concentrations of GlcNAc (50 mM and 100 mM) further enhanced hyphal formation, whereas 30 mM GlcNAc did not induce filamentation. A custom-designed experiment using JMP software was employed to assess the effects of GlcNAc and MgSO₄ combinations, identifying the optimal conditions as 30 mM GlcNAc and 0.05 mM MgSO₄, with an overall desirability score of D = 0.99. Run 5, 7, 9, 10, 11, 12, 13, 17, 18, 19, and 20 also induced hyphal formation. Statistical analysis indicated that GlcNAc was the primary contributing factor (*p* < 0.05, R² = 0.26, *p* = 0.043), whereas MgSO₄ was not significant. Furthermore, gene expression analysis revealed the transcriptional upregulation of hyphal-specific genes such as *HWP1* and *HGC1*, suggesting that hyphal induction could be driven by GlcNAc alone or in combination with nutrient-deprived stress responses. pH analysis during hyphal formation showed an initial shift toward alkalinity at the 1 hour and 2 hours time points, followed by lowering to acidic pH at 4 hours and 6 hours, coinciding with hyphal formation. The hyphal induction achieved under these optimized conditions may provide insights in into morphogenic transitions associated with pathogenic Candida strains. This study holds potential to enhance our understanding of the molecular mechanisms underlying virulence traits such as hyphal formation, both *in-*vitro and *in*-vivo. Such insights could aid in the development of host-pathogen interaction models, facilitate antifungal drug target discovery, and contribute to the identification of Candida-specific pathogen associated molecular patterns (PAMPs) that elicit damage associated molecular patterns (DAMPs) mediumted host responses. This optimized medium may serve as a useful tool in guiding the development of targeted and preventive therapeutic strategies against Candidiasis.

## CRediT authorship contribution statement

**Rachana Arvind:** Writing – original draft, Methodology, Investigation, Conceptualization.

**Atheena PV:** Visualization, Investigation, Formal analysis.

**Allan Britto J**: Methodology, Investigation.

**Uttara Chakraborty:** Writing – review & editing, Supervision, Data curation.

**Ritu Raval:** Writing – review & editing, Supervision, Project administration, Data curation.

## Data availability

Data will be made available on request.

